# A simple chromogenic whole-cell arsenic biosensor based on *Bacillus subtilis*

**DOI:** 10.1101/395178

**Authors:** Niels Wicke, David S. Radford, Christopher E. French

**Affiliations:** School of Biological Sciences, University of Edinburgh, Roger Land Building, Alexander Crum Brown Road, Edinburgh EH9 3FF, UK

## Abstract

Arsenic contaminated ground water is a serious public health issue, and recent estimates place 150 million people worldwide at risk. Current chemical field test kits do not reliably detect arsenic at the lower end of the relevant range, and may generate toxic intermediates and waste. Whole-cell biosensors potentially provide an inexpensive, robust and analyte-specific solution to this problem. The second generation of a *Bacillus subtilis*-based arsenic biosensor, designated Bacillosensor-II, was constructed using the native chromosomal *ars* promoter, *arsR* and the reporter gene *xylE* encoding catechol-2,3-dioxygenase. Within four hours, Bacillosensor-II can detect arsenic in the form of arsenate AsO_4_^3-^ at levels more than one order of magnitude below the recommended safe limit for drinking water suggested by the World Health Organisation (10 µg/L). Detection is reported by the enzymatic conversion of the inexpensive substrate catechol to 2-hydroxy-*cis,cis*-muconic semialdehyde, a bright yellow product visible by eye. We hope that this work will aid in developing a simple inexpensive field test kit for screening of drinking water for arsenic contamination.

## Introduction

Arsenicosis is a chronic health condition characterized by cutaneous lesions triggered by prolonged ingestion of the trace element arsenic (As), predominantly as arsenate AsO_4_^3-^ and the more toxic form arsenite AsO_3_^3-^. Progression manifests in peripheral vascular disease and cancers of internal organs (Das and Sengupta, 2008, Kay, 2011). Arsenic is mobilized from the earth’s crust by geogenic leaching processes and consequently contaminates groundwater (Smedley and Kinniburgh, 2002). Arsenic may also accumulate in rice (*Oryza sativa*) when irrigated with contaminated groundwater (Zhao et al, 2010), thereby presenting a major and perhaps under-appreciated route of ingestion. The safe limit for As in drinking water recommended by the World Health Organisation (WHO) is 10 µg/L (ppb), though many governments in affected areas, like Bangladesh, place this limit at 50 µg/L (Flanagan et al., 2012). Other affected areas include Nepal (Panthi et al., 2006) and Vietnam (Trang et al., 2005). It is currently estimated that 150 million people are at risk worldwide; however, a recent study in Pakistan suggests this figure may need to be revised upward (Podgorski et al., 2017).

Given the remote locality and economic situation of affected areas, there is an urgent need for field-test kits that are inexpensive, simple to operate and interpret, do not require a cold-chain or specialized equipment, and do not generate toxic waste, whilst being highly specific to bio-available forms of As (detecting both arsenate and arsenite) and able to reliably detect below the recommended limit of 10 ppb. Commercially available chemical field kits are based on the Gutzeit method, which generates toxic arsine gas (AsH_3_) by reacting water samples with metallic zinc and hydrochloric acid. A coloured spot is then produced by reacting AsH_3_ with indicator paper coated in mercuric bromide where the intensity of the spot indicates the concentration of As in the water sample. Presently available kits (Panthi et al, 2006, Safarzdeh-Amiri et al., 2011, Deshpande et al., 2001) suffer from inadequate sensitivity, generation of false negatives at low concentrations, and inaccuracy due to human interpretation of the visual output. Inclusion of a colorimetric instrument to ameliorate variability, for example in the Wagtech Digital Arsenator (Safarzdeh-Amiri et al., 2011), raises the initial investment and operating cost. Finally, AsH_3_ is toxic and the mercuric salts present a disposal issue (French et al., 2012).

Biological systems are composed of highly specific and sensitive molecular recognition systems. Biosensors couple biological sensing entities to a transduction element that communicates the recognition event, for example through luminescence or electrical current, in a quantitative manner. Various types of biosensor able to detect arsenate and arsenite have been reported in the literature (reviewed by French et al., 2012). Enzymic biosensors use enzymes as recognition elements. Redox enzymes are particularly suitable because they generate an electrical current upon interaction with their substrate/analyte. For example, Male et al. (2007) immobilized arsenite oxidase from *Rhizobium* sp. NT-26 on a carbon electrode enabling As detection with a linear response between 1 and 500 µg/L. Enzymes, however, may be expensive to produce and are more or less labile, which may pose challenges in distribution and storage. Alternatively, whole cell biosensors, also known as bioreporters, use living cells as the recognition element. Whole cells have the advantage of being self-manufacturing and sustaining, can integrate detection of multiple analytes, and can be lyophilized for distribution and storage. Whole-cell arsenic biosensors are generally genetically modified bacteria which couple induction of the arsenic-responsive *ars* promoter to expression of a reporter gene which gives a quantifiable output. The *ars* promoter is controlled by the repressor ArsR, which in the presence of arsenite, dissociates from the promoter allowing transcription. Whole cell arsenic biosensors have generally been based on the *ars* promoters of *Escherichia coli* (Diorio et al., 1995), *Staphyloccocus* plasmid pI258 (Ji and Silver, 1995) and *Bacillus subtilis* 168 *skin* element (Sato and Kobayashi, 1998), with reporter genes such as *lacZ* (β-galactosidase), bacterial or firefly luciferase, or Green Fluorescent Protein (GFP). For example, Ji and Silver (1992) combined the *ars* promoter and *arsR* from *Staphylococcus* pI258 with *lacZ* and *luxAB* in *Staphylococcus aureus* and *E. coli*. Both host strains could detect 0.1 µM arsenite (7.5 µg/L As). With awareness of the dangers of arsenic contaminated groundwater growing, a few arsenic biosensors have undergone field trials. Notably, an *E. coli* sensor featuring the pI258 *ars* promoter and red-shifted GFP performed similarly to chemical tests of As-contaminated water in Yun-Lin County Taiwan (Liao et al., 2005). For a comprehensive review of arsenic biosensors, see French et al. (2012) and Diesel et al. (2009). However, regulations regarding the use of genetically modified microorganisms outside the laboratory hamper the commercial development of such devices.

Previously, two arsenic biosensors have been developed in this laboratory. The first is a pH-based system that exploits the ‘mixed-acid’ fermentation of *E. coli* in the presence of lactose, leading to acid formation in the presence of arsenic, easily detectable using a simple pH indicator such as bromothymol blue (de Mora et al, 2011). The second, termed Bacillosensor-I, uses the endospore-forming Gram-positive bacterium *B*. *subtilis* as host chassis. Endospores are dormant highly resistant forms capable of surviving in harsh physiochemical conditions for centuries. The potential advantages for storage and distribution are obvious. Like *E. coli, B. subtilis* grows at temperatures up to 45°C allowing its use in tropical countries. Bacillosensor-I was composed of the chromosomal *B. subtilis ars* promoter and *arsR* fused to the reporter gene *xylE* on plasmid pVK168, but was not sufficiently sensitive to arsenic concentrations below 75 ppb (µg/L)(Joshi et al, 2009). Here we describe a modified version, Bacillosensor-II, with enhanced sensitivity and responsiveness.

## Results

Bacillosensor-II was assembled using BioBricks (Knight, 2003). As in Bacillosensor-I (Joshi et al., 2009), the genetic sensing and output elements consist of the *B. subtilis ars* promoter and its repressor (P_*ars*_*-arsR*, BBa_J33206) fused to the reporter gene *xylE* (BBa_J33204) from *Pseudomonas putida* pWW0. XylE (catechol-2,3-dioxygenase) catalyses the oxidation of catechol, a cheap colourless substrate, to the bright yellow compound 2-hydroxy-*cis,cis*-muconic semialdehyde (HCMS) which absorbs strongly at 377 nm. Bacillosensor-II was assembled in a BioBrick-compatible form of pTG262 (BBa_I742123), a shuttle vector with a Gram-positive origin of replication compatible with *Escherichia coli* (Shearman et al., 1989). The construct was assembled in *Escherichia coli* and then transformed into *B. subtilis* 168.

To investigate the response of the Bacillosensor to arsenate, cells were induced overnight in the presence of different concentrations of arsenic (as sodium arsenate) prior to addition of catechol for 1 hour to allow colour development (Figure 1). Significant induction of XylE activity is evident in 5 ppb arsenate and colour intensity correlates positively with increasing arsenate concentration.

**Figure 1.**
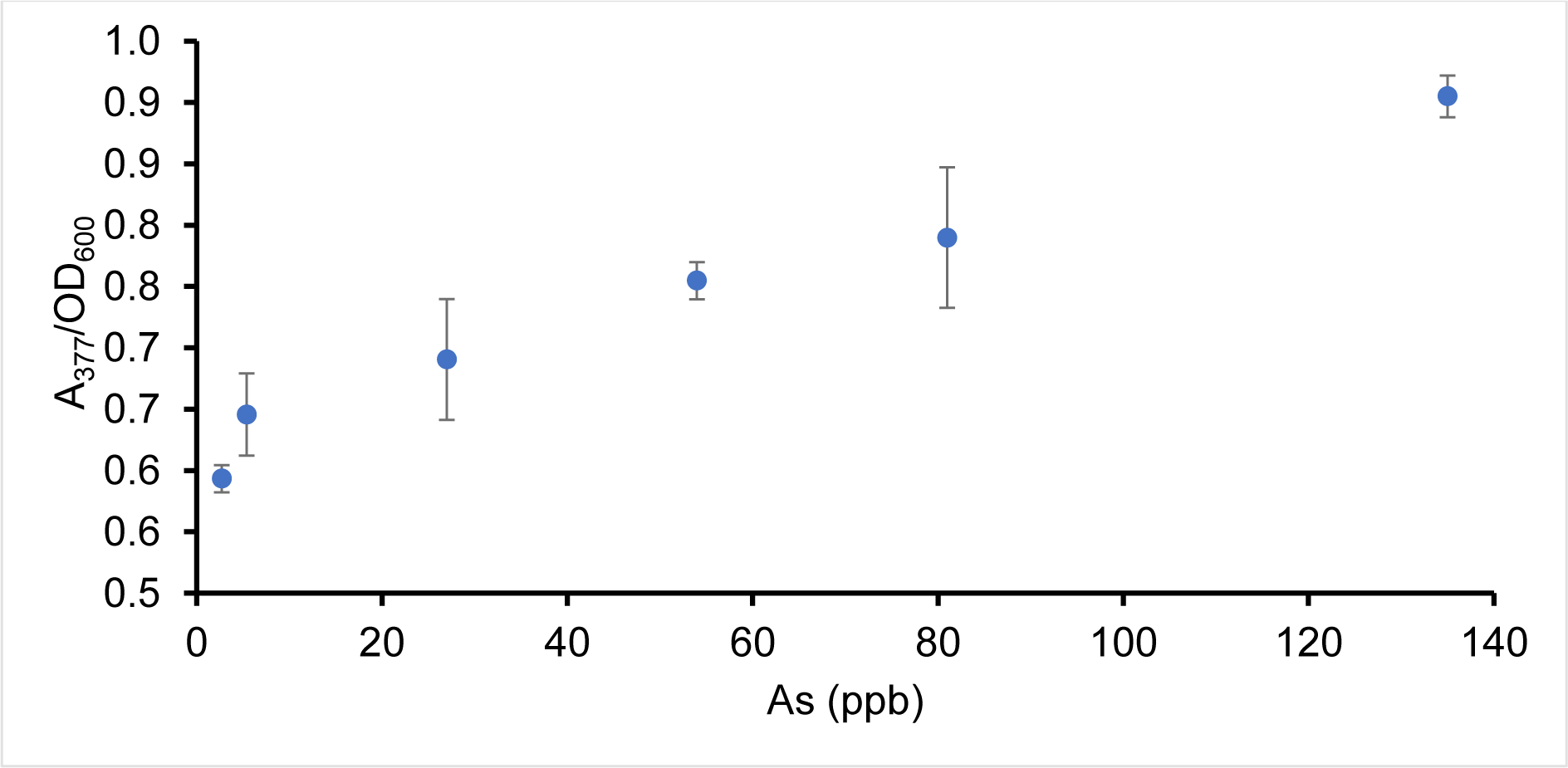
Average A_377_/OD_600_ of supernatant of cultures grown overnight in presence of arsenate prior to incubation with 0.5mM catechol for 1 hour at room temperature. Average A_377_/OD_600_ of cultures grown in the absence of arsenate were subtracted. Lack of XylE activity in wild-type *B. subtilis* was shown previously (Joshi et al, 2009) and is thus omitted here. Errors bars indicate standard deviation of technical triplicates.

Another desirable property of (field) biosensors is a rapid response time. This includes the time required for induction and enzyme production, as well as the time required for colour development. The latter was tested by growing an overnight culture in the presence of various concentrations of arsenate and monitoring the increase in HCMS over time following addition of catechol (Figure 2). It is assumed that *xylE* is maximally induced, relative to the concentration of arsenate, following overnight incubation. At concentrations of 50 ppb arsenate and above the presence of HCMS is visible by eye in less than one minute (within the time taken from addition of catechol to making the first measurement). At such levels of arsenate, colour development appears to plateau after 60 minutes. In the lowest arsenate concentration tested, 0.54 ppb As (1 ppb arsenate), the bright yellow colour of HCMS was clearly visible 2 hours after catechol addition.

**Figure 2.**
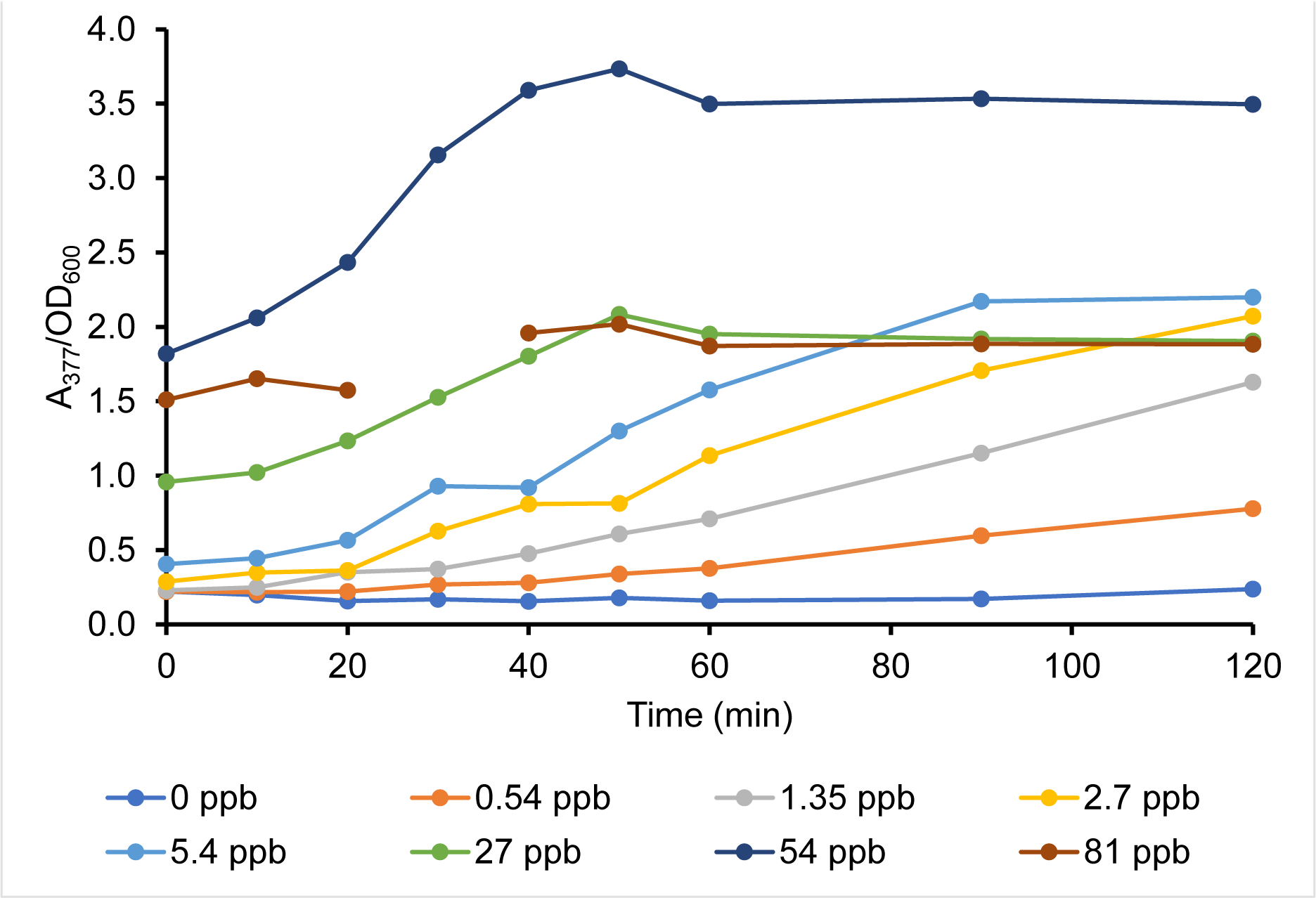
Conversion of catechol to HCMS over time in Bacillosensor-II cultures induced overnight with arsenate. Cultures were grown overnight in presence of arsenate. Catechol was added to 0.5mM and aliquots taken periodically and the A_377_ of the supernatant measured immediately. Measurement for 81 ppb at 30 min is missing due to human error.

**Figure 3.**
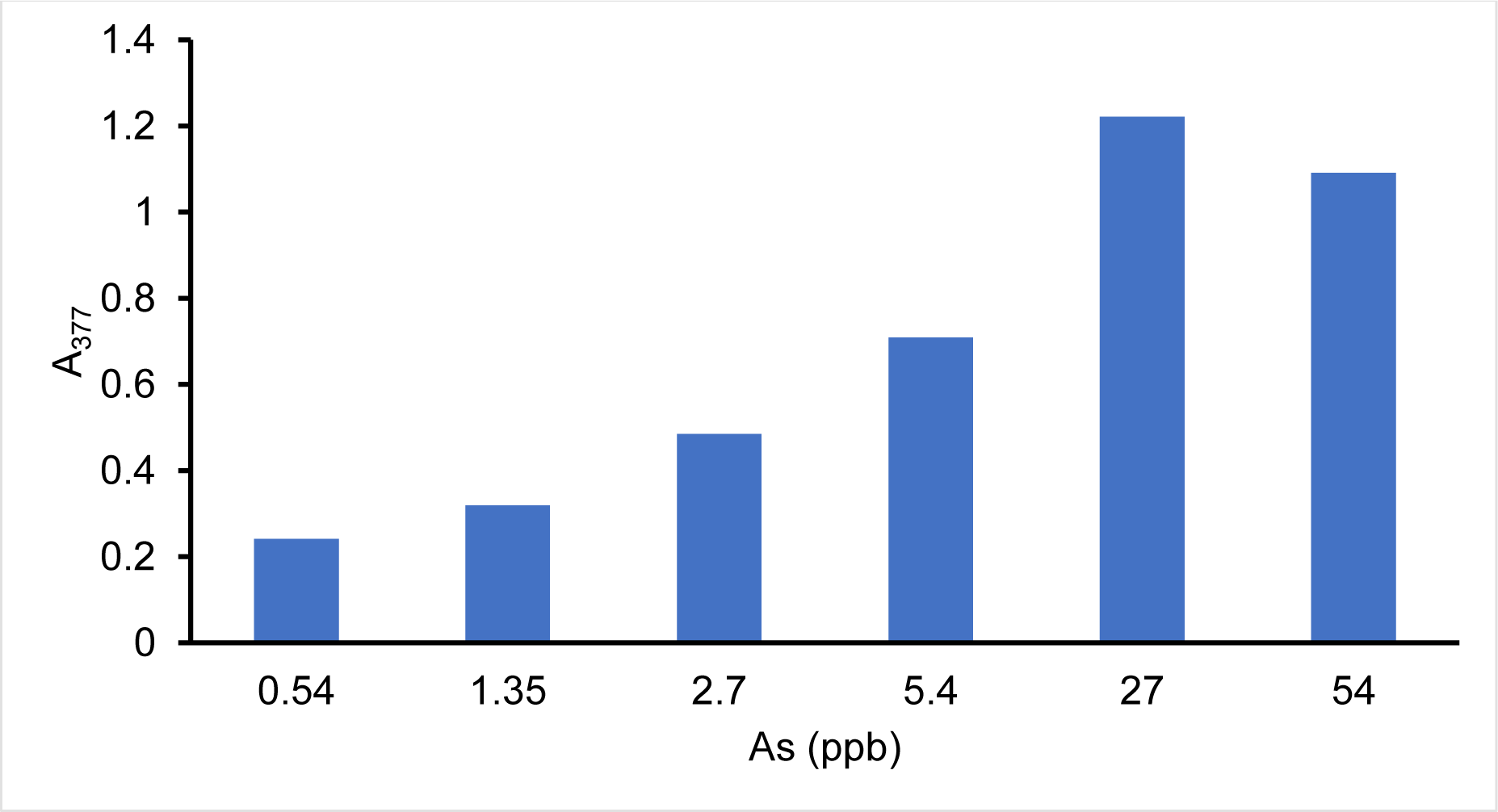
A_377_ of Bacillosensor-II culture supernatant after incubation with 0.5mM catechol and 0, 1, 2.5, 5, 10, 50 or 100 ppb arsenate for 4 hours at 37°C with shaking at 200 rpm. A sample with catechol but no arsenate was used as blank.

A potential complication of this system is that in solution and in the presence of oxygen catechol is oxidized to a brown melanin-like polymeric substance which obscures the yellow colour of HCMS. Oxygen cannot be excluded, since both *B. subtilis* growth and XylE activity are O_2_-dependent. Thus, catechol cannot be included in the medium prior to overnight incubation. An experiment was conducted to determine whether XylE production and activity can be observed prior to degradation of catechol, when catechol and arsenate are added simultaneously (Figure 3). Results showed that Bacillosensor-II can detect 0.54 ppb As as arsenate within four hours after simultaneous addition of arsenate and catechol. Other experiments indicated that the time taken for non-enzymic degradation of catechol is strongly dependent on the incubation conditions. In our hands, catechol in 5 ml culture medium in a sealed 50 ml centrifuge tube remained stable for more than 6 hours (data not shown). A representative image of colour development of overnight-arsenate-induced cultures incubated with catechol for 1 hour is shown in Figure 4.

**Figure 3.**
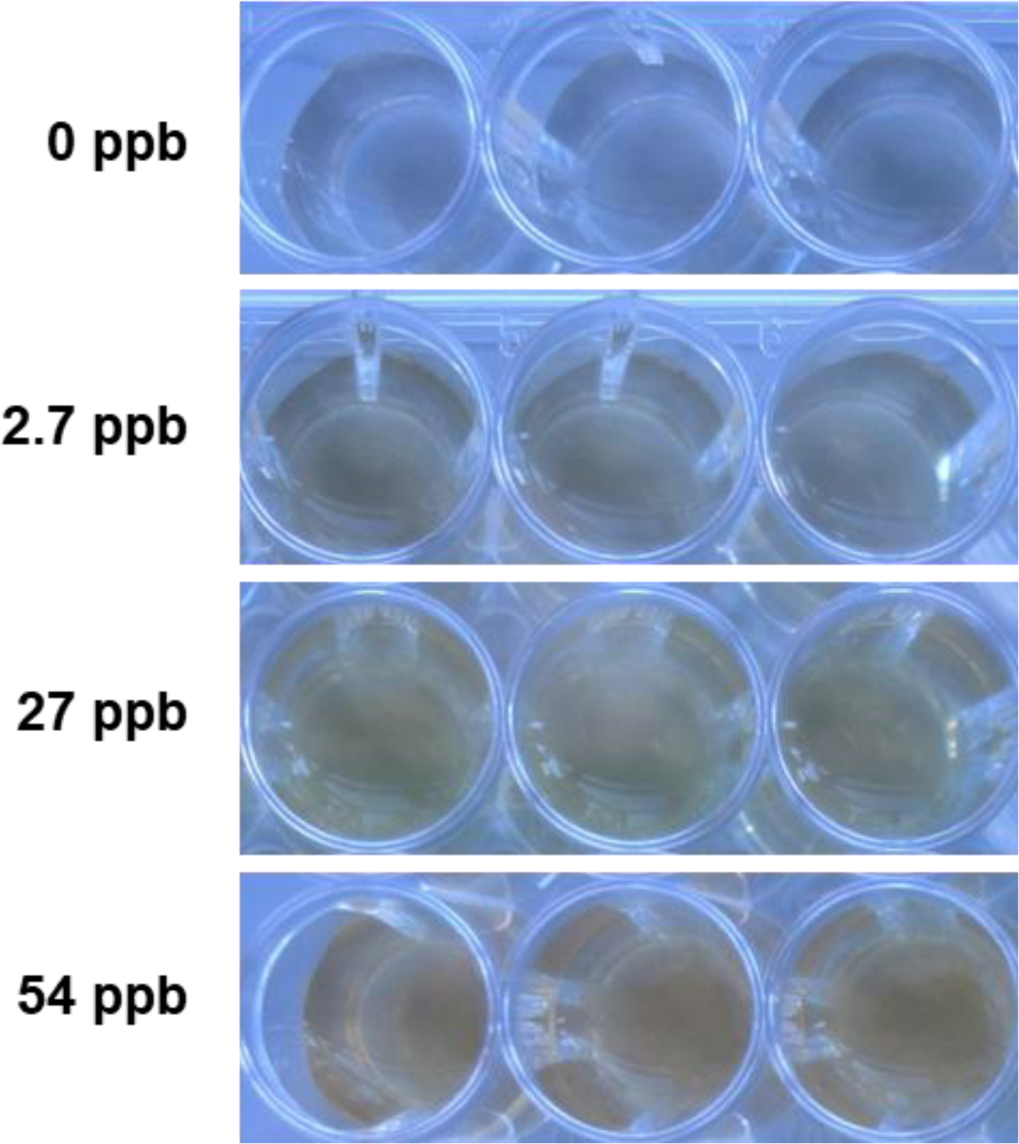
Representative image of the yellow colour of HCMS. Cultures were grown overnight in presence of arsenate. Catechol was added to 0.5 mM and incubated for 1 hour at room temperature. Cells were removed by centrifugation. Note the slight brown colour in samples with 2.7 ppb, which may represent oxidation of catechol as well as conversion to HCMS.

GM regulatory issues are very important in moving towards commercialization of genetically modified whole cell biosensors. As a step towards generating a system with no antibiotic selection marker, we used our newly developed BacilloFlex DNA assembly system (Wicke et al., 2017) to generate a version of Bacillosensor-II in which the *P*_*ars*_-*xylE* cassette (omitting *arsR*, and relying on the chromosomal *arsR* gene for ArsR synthesis) is integrated on the genome at the *amyE* locus. In principle this would allow subsequent removal of the chloramphenicol resistance gene. The new system, Bacillosensor-III, was tested with arsenate in the same way as Bacillosensor-II, and showed comparable sensitivity, with about half the colour intensity of the multi-copy plasmid system (Figure 5).

**Figure 5.**
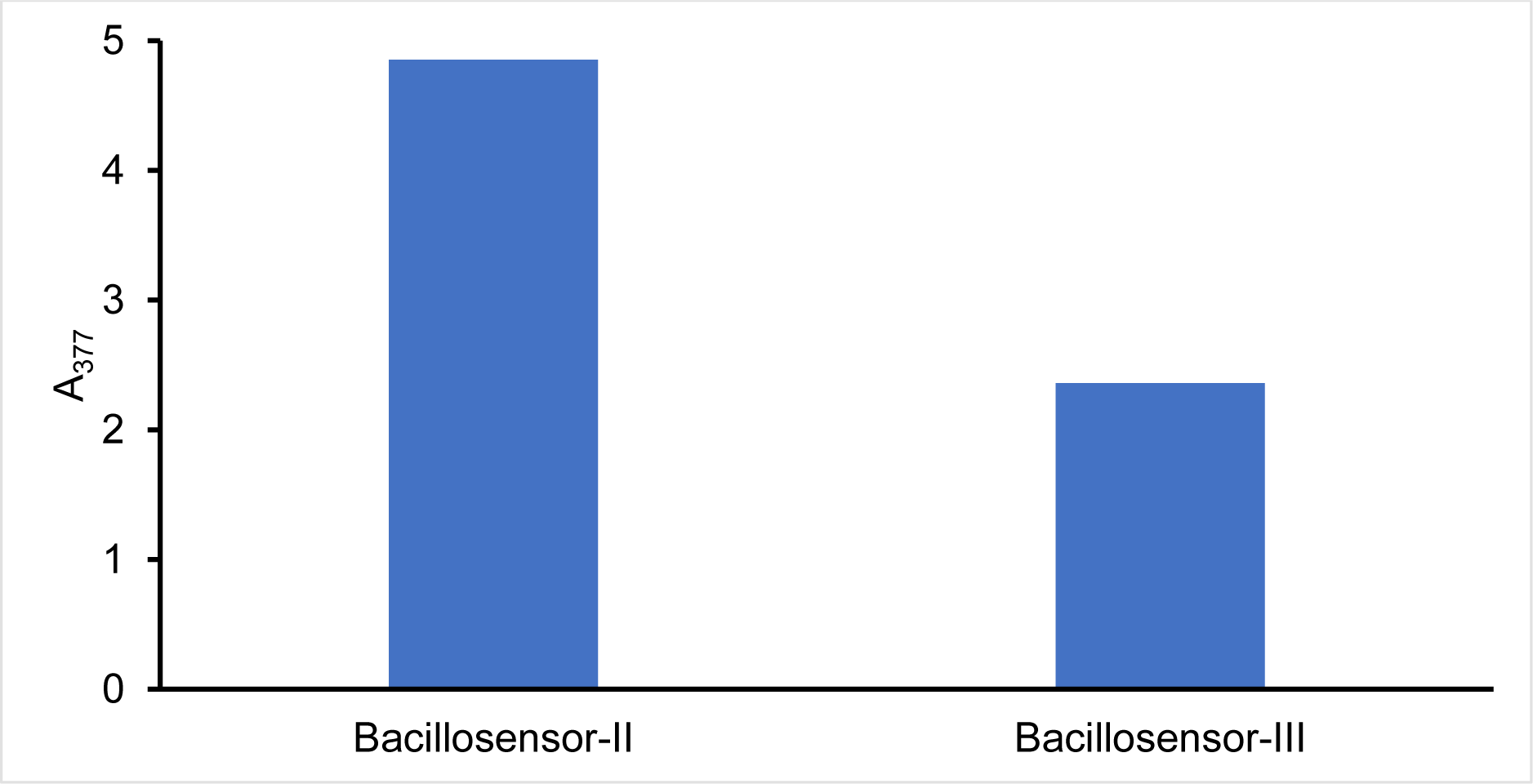
XylE activity of Bacillosensor-II and -III. Cultures were grown overnight with 10 ppb arsenate (5.4 ppb As) and incubated with 0.5 mM catechol for 6 hours. A_377_ was blanked against supernatant from a wild-type *B. subtilis* control.

## Discussion

The aim of this work was to construct and test a *B. subtilis-*based arsenic biosensor suitable for field applications. The data in Figure 1 demonstrate the sensitivity of Bacillosensor-II to 5 ppb arsenate (2.7 ppb As) which is below the limit for drinking water recommended by the WHO. It is worth noting that some arsenate toxicity was reflected in the final optical densities of overnight cultures. Although a positive correlation between arsenate and HCMS concentration was observed, further experiments are needed to determine whether this system is suitable for accurately quantifying the levels of bio-available arsenate, or whether it can only report on the presence (above a minimal threshold) or absence of arsenic.

Data in Figures 2 and 3 demonstrate the short time-frame within which a response can be obtained, an important parameter in the field. It is also important that the output substrate and sample/analyte may be added simultaneously which contributes to the ease of performing the test. Our results suggest this is possible, despite the instability of catechol; however, careful design of the final sensor device would be required to minimize non-enzymic catechol degradation. Our results also indicate that the biosensor cassette can be integrated onto the chromosome, allowing for later removal of the antibiotic resistance gene, which may facilitate regulatory approval.

Taken together, this work is encouraging progress toward a whole-cell *B. subtilis* arsenic biosensor sensitive to levels of arsenic below the recommended limit for drinking water by the WHO. This system is simple to operate at ambient temperatures, utilizes the inexpensive substrate catechol, is self-manufacturing and maintaining, and has the potential to excel at storage and transport due to the inherent stability of *Bacillus* endospores. The output is visible by eye and does not require expensive equipment.

Additional work should therefore include preparing and testing endospores of the Bacillosensor-II strain. Ideally, these could be germinated directly in growth medium with added water sample and catechol to give a rapid read-out. In addition, experiments should be conducted to test the response to arsenite; generally such systems are more strongly induced by arsenite than arsenate (Sato and Kobayashi, 1998), since arsenate must be reduced to arsenite by arsenate reductase (ArsC) prior to interacting with ArsR.

In addition to our own Bacillosensor-I (Joshi et al, 2009), previously described arsenic biosensors based on *B. subtilis* include those of Tauriainen et al. (1997), using firefly luciferase as reporter, and Date et al. (2007), using either GFP or a chemiluminescent system based on β-galactosidase. In the latter case it was reported that spores could be stored in dry form for more than 6 months prior to use. Provided that the issues of catechol stability can be overcome, XylE as reporter offers certain potential advantages, in that the substrate is very cheap, and the reaction product can be easily detected by eye.

To the best of our knowledge, no whole-cell arsenic biosensors are commercially available despite their potential superiority over current chemical field kits. We hope this work will contribute to providing an inexpensive widely-adopted system for detection of arsenic in drinking water. A chromogenic biosensor based on *Bacillus* endospores has many advantages, and development of such a device is also the focus of the Arsenic Biosensor Collaboration (www.arsenicbiosensor.org).

## Methods

### Strain and culture conditions

Routine growth of *Bacillus subtilis* 168 was carried out on Luria Bertani (LB) agar or broth at 37°C (shaking at 200 rpm for liquid cultures) with 10 µg/ml chloramphenicol (Acros Organics)(cml10) as necessary.

### Materials

Catechol and sodium arsenate heptahydrate were purchased from Sigma-Aldrich and stocks of 0.5M and 10 000 ppb (or 10 µg/ml arsenate, 22.4 µg/ml sodium arsenate heptahydrate in sterile water) were prepared, respectively.

### Generation of Bacillosensor II and III

The parts BBa_J33206 (P_*ars*_ -*arsR*) and BBa_J33204 (*xylE* reporter) were inserted into BBa_I742123 (pTG262) by the standard RFC10 BioBrick protocol (Knight, 2003) and transformed into *B. subtilis.* Competent cells were prepared, stored and transformed by the Groningen Method (Bron, 1990). Bacillosensor-III was generated using parts and methods described in Wicke et al. (2017) to introduce *P*_*ars*_-*xylE*, together with a chloramphenicol resistance marker, onto the chromosome at the *amyE* locus.

### Measuring concentration of HCMS

The concentration of HCMS was measured by absorbance at 377 nm in a SpectroSTAR Nano (BMG Labtech). Measurements were made using culture supernatants to avoid interference due to light scattering by cells. Samples were diluted 1/5 where necessary. To correct for cell density, OD_600_ of cultures was also recorded.

### Testing responsiveness of Bacillosensor-II to arsenate

Arsenic assays were performed by inoculating 5 ml of LB + chloramphenicol medium in 50 ml disposable centrifuge tubes with 10 µl of overnight culture, with addition of sodium arsenate to 0, 5, 10, 50, 100, 150 or 250 ppb (corresponding arsenic concentrations are indicated in Figure 1). Cultures were grown for 19 hours at 37°C with shaking at 200 rpm. Triplicate 1 ml samples were transferred to clean sterile microfuge tubes and catechol added to 0.5 mM. Samples were incubated at room temperature with an open lid for 1 h before centrifugation and transfer of supernatant to a 48-well plate (Greiner Bio-One) for measurement of A377.

### Testing response time of cultures after catechol addition following overnight induction

To determine the time required for colour development, a similar experiment was performed, but at each time point 100 µl of sample was transferred to a microcentrifuge tube and cells removed by centrifugation. Supernatant (90 µl) was transferred to a 96-well plate (Greiner Bio-One) and A_377_ was measured immediately. A_377_ was divided by OD_600_ of the cultures to correct for differences in cell density.

### Testing responsiveness to simultaneous addition of arsenate and catechol

A 10 ml Bacillosensor culture was grown overnight, and samples of 1 ml transferred into 15 ml centrifuge tubes and arsenate added to 0, 1, 2.5, 5, 10, 50, 100 or 150 ppb. Assays were incubated at 37°C with shaking at 200 rpm for 4 h. Cells were removed by centrifugation and 0.9 ml supernatant transferred to plastic cuvettes for measurement of A_377_.

### Testing responsiveness of Bacillosensor-III to arsenate

Bacillosensor-III was grown in LB broth with 10 µg/ml chloramophenicol, incubating at 37°C with shaking at 200 rpm. Arsenic assays were performed by inoculating 5 ml of the same medium in 50 ml disposable centrifuge tubes with 10 µl of overnight culture, with addition of sodium arsenate to 10 ppb (5.4 ppb As). Cultures were grown for 18 h at 37°C with shaking at 200 rpm. Catechol was added to a final concentration of 0.5 mM., and incubation continued for a further 6 h. Cells were removed by centrifugation and 900 µl supernatant transferred to a plastic cuvette. A wild-type *B. subtilis* 168 sample was used as blank.

